# Molecular mechanisms for environmentally induced plasticity in the positioning of meiotic recombination at hotspots

**DOI:** 10.1101/2020.09.24.312371

**Authors:** Tresor O. Mukiza, Reine U. Protacio, Mari K. Davidson, Wayne P. Wahls

## Abstract

In meiosis, Spo11/Rec12-initiated homologous recombination is clustered at hotspots that regulate its frequency and distribution across the genome. Intriguingly, the intensities and positions of recombination hotspots can change dramatically in response to intracellular and extracellular conditions, and can display epigenetic memory. Here, using the fission yeast *Schizosaccharomyces pombe*, we reveal mechanisms for hotspot plasticity. We show that each of six hotspot-activating proteins (transcription factors Atf1, Pcr1, Php2, Php3, Php5, Rst2) is rate-limiting for promoting recombination at its own DNA binding site, allowing each class of hotspot to be regulated independently by agonistic and antagonistic signals. We also discovered that the regulatory protein-DNA complexes can establish a recombinationally poised epigenetic state before meiosis. Notably, Atf1 and Pcr1 controlled the activation of DNA sequence-dependent hotspots to which they do not bind; and they do so by regulating the expression of other hotspot-activating proteins. Thus, while each transcription factor activates its own class of DNA sequence-dependent hotspots directly in *cis*, cross-talk between regulatory networks modulates in *trans* the frequency and positioning of recombination at other classes of DNA sequence-dependent hotspots. We posit that such mechanisms allow cells to alter the frequency distribution of meiotic recombination in response to metabolic states and environmental cues.

## Introduction

In meiosis cells express the broadly conserved Spo11/Rec12 protein which, along with other components of the basal meiotic recombination machinery, catalyzes the formation of dsDNA breaks (DSBs) that initiate homologous recombination (1-3). The broken chromosome uses its homologous partner for template-directed repair, which yields recombinant chromosomes that harbor detectable gene conversion events, a subset of which have associated crossing over. Consequently, at any given location in the genome, there is general correspondence between the frequency of DSBs and the frequency of recombinants. In many, diverse taxa (e.g., fungi, plants, vertebrates), the DSBs and recombination events cluster at “hotspots” that regulate the frequency and distribution of recombination across the genome (4-7).

In a subset of metazoans (e.g., mice, cattle, primates), the positions of hotspots are shaped by Prdm9, and changes in its DNA binding domain confer dramatic, evolutionarily rapid changes in the distribution of recombination (8,9). Many other, sequence-specific DNA binding proteins that do not evolve as rapidly also position recombination at hotspots (10) and, correspondingly, hotspot locations are more stable in species that lack Prdm9 (e.g., birds, amphibians, many fishes, canids, marsupials, plants, fungi). The global distribution of DSB hotspots can be defined with precision using several molecular tools (11-17); and the reproducibility of such measures is exemplified by two, extensively studied, highly diverged organisms, fission yeast and budding yeast. Experiments conducted using different detection methods, at different times, and even in different laboratories have yielded similar patterns for the distribution of DSB hotspots within each of these species, differing primarily in the resolution and sensitivity of the assay employed (14-20). However, most of the intra-species comparative studies have used similar genetic backgrounds and experimental conditions, such as the media and temperature in which meiosis was induced. Remarkably, other experiments revealed that differences in metabolic states and environmental cues can trigger dramatic changes in the positioning of recombination at hotspots, even within isogenic or genetically identical strains.

Plasticity in the frequency distribution of meiotic recombination is a long-recognized, well-documented phenomenon. For example, differences in sex or mating type affect the distribution of recombination among hotspots of diverse species (e.g., fungi, mammals), as does the addition of an ectopic mating type cassette (21-24). Both auxotrophies and nutritional states, such as the addition or removal of amino acids in the media, affect hotspot activity (25,26). Differences in parental mating type and the freezing of diploids each affect the activity of hotspots in subsequent meiosis, which suggests that there is epigenetic imprinting (21,27). Such imprinting can also be inferred from the structure and composition of chromatin at hotspots (28-30). Differences in temperature during meiosis affect patterns of recombination across the genomes of diverse species (31), and in two of these species the effects of temperature on the global distribution of recombination-initiating DSBs have been examined. In budding yeast only about 20% of DSB hotspots, as defined using a frequency threshold, occur at the same positions when meiosis is carried out at 14 °C, 30 °C and 37 °C (32). In fission yeast differences in temperature affect the distribution of DSBs (23), as well as rates of recombination (33), at DNA sequence motif-dependent hotspots (34,35). Hypothetically, such regulatory DNA sites and their binding proteins might contribute to the plasticity of recombination positioning (30,32). There are other, even more perplexing manifestations of plasticity. For example, while Prdm9 can modulate the initiation of recombination at its own DNA binding sites (24), the deletion of Prdm9 leads to the generation of new hotspots elsewhere in the genome (36). A similar situation applies for transcription factors Bas1 and Ino4 of budding yeast, whose removal represses DSBs at some hotspots and induces DSBs at others (37,38). In summary, environmental conditions and metabolic states can reshape—in some cases quite dramatically and by yet unknown mechanisms—the frequency distribution of meiotic recombination across the genomes of diverse taxa.

To gain insight into potential mechanisms for the plasticity of recombination events, we studied three exceptionally well characterized classes of recombination hotspots in fission yeast. Each class of hotspot is regulated by a discrete DNA sequence motif that has been defined functionally at single-nucleotide resolution by scanning, base pair substitutions in the genome (**Table 1**) (39-41). In each case, the binding of transcription factors to those DNA sites is essential for hotspot activity. The Atf1-Pcr1 heterodimer (of the ATF/CREB/AP-1 family) promotes recombination at *M26* (*CRE*-like) DNA sites (5,34,42); Php2-Php3-Php5 complex activates recombination at *CCAAT* box DNA sites (41); and Rst2 regulates recombination at *Oligo-C* DNA sites (41). These disparate *cis*-acting regulatory modules each share a common downstream mechanism. Each protein-DNA complex triggers the displacement of nucleosomes to promote access of the basal recombination machinery (Spo11/Rec12 complex) to its DNA substrates within chromatin (30), thereby stimulating locally the frequency of recombination-initiating DSBs (5,43). Here we describe molecular mechanisms by which these *cis*-acting regulatory modules can reshape the frequency distribution of recombination across the genome in response to metabolic and environmental cues.

**Table 1.**
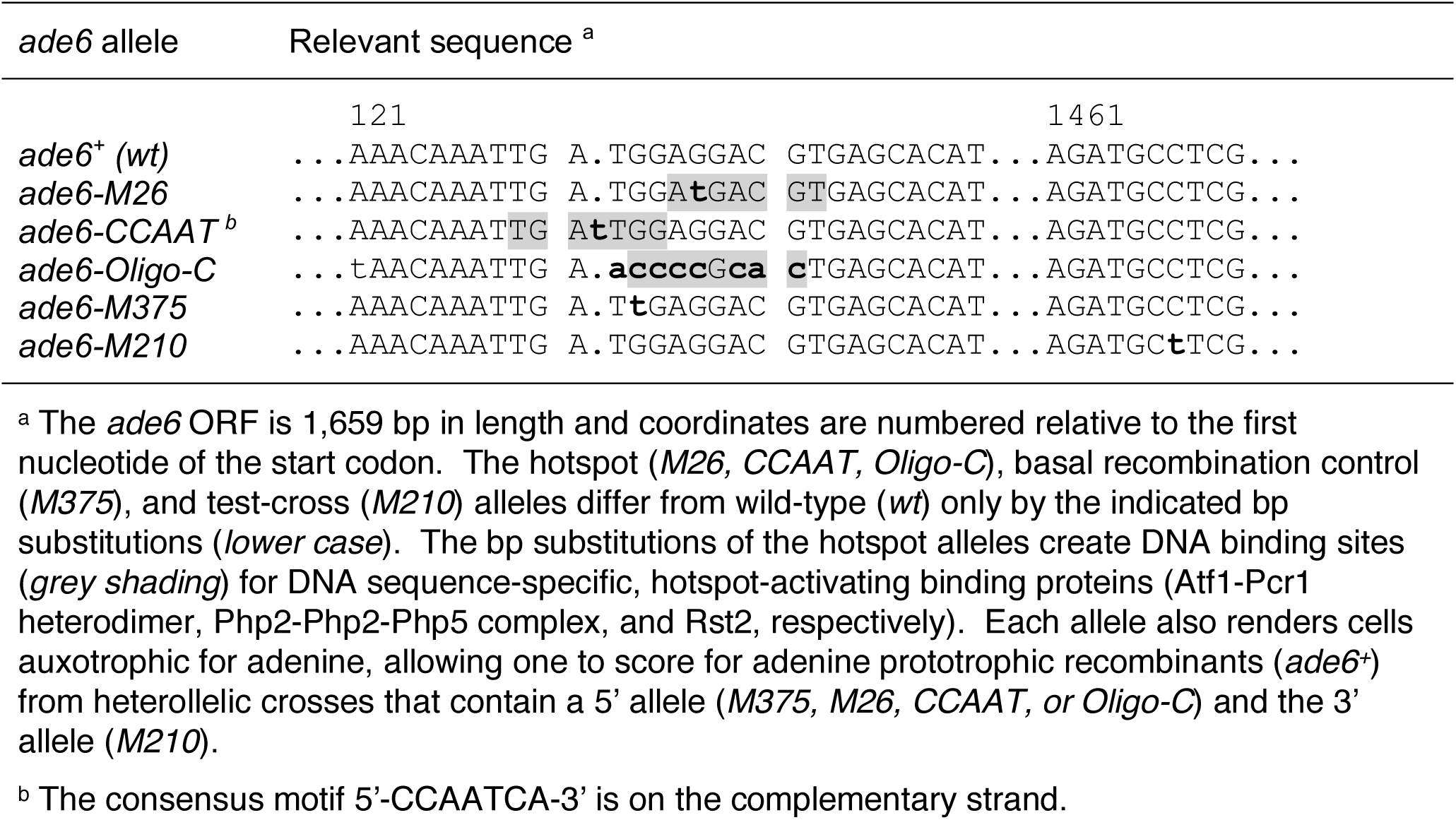
DNA sequences of *ade6* alleles.

## Materials and Methods

### Yeast culture and strain construction

The DNA sequences of *ade6* alleles used in this study are provided in **Table 1** and the genotypes of all strains are listed in **Table 2**. Strains were constructed using standard genetic techniques and were cultured in rich or minimal media supplemented as necessary with amino acids and bases at 100 µg/ml (44-46).

**Table 2.**
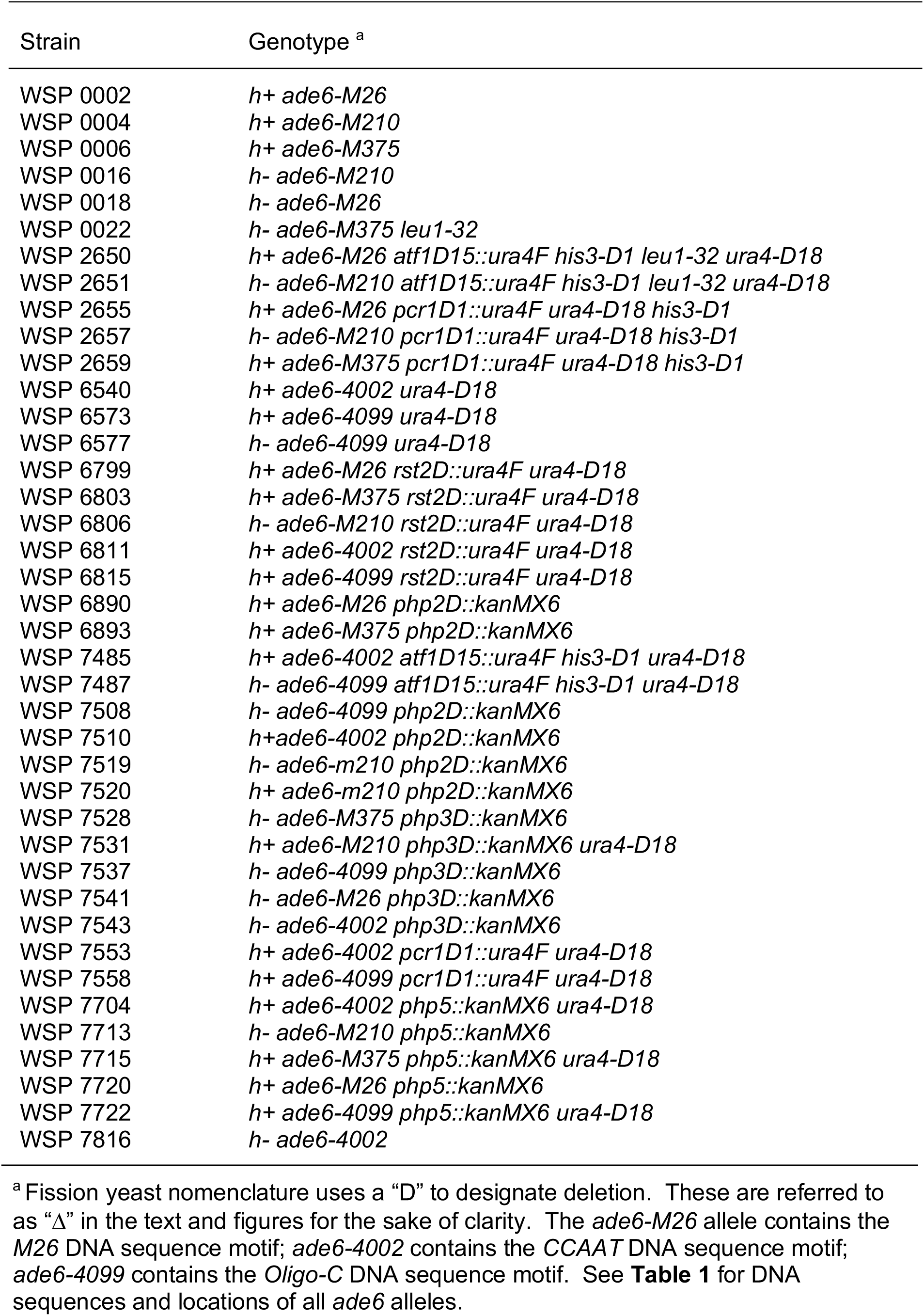
Genotypes of fission yeast strains. [upon acceptance for publication, this table will be moved to a supplementary online materials file]

### Measurements of meiotic recombination

Methods to measure rates of meiotic recombination were as described (34,47). In brief, haploid strains with different *ade6* alleles (**Table 1**) were mated; then, haploid meiotic products (spores) were harvested, and serial dilutions of spores were plated on minimal media that contained or lacked adenine. The titer of Ade^+^ recombinant spore colonies was divided by the titer of all viable spore colonies to yield the recombinant frequency of each cross. Frequencies reported in the figures are mean ± SEM of values from three or more independent biological replicates of each cross.

### Measurements of mRNA abundance

Quantitative, real-time, reverse-transcription PCR (qRT-PCR) was used to measure the abundance of mRNAs in cells (A595 = 0.5) that had been starved for nitrogen for 60 minutes. For each sample, total RNA from approximately 1 × 10^8^ cells was extracted using the Quick-RNA Fungal/Bacterial Miniprep (Zymo Research, Irvine, CA). For each sample, 1 µg of RNA was treated with TURBO DNase using the TURBO DNA-*free*™ Kit, which includes reagents for the digestion of DNA along with the removal of the enzyme and divalent cations post-digestion. Following DNase treatment, cDNA was synthesized from 400 ng RNA as template using the iScript cDNA Synthesis kit (BioRad, Hercules, CA). The cDNA was used as template for qPCR using BlazeTaq SYBR Green qPCR Master Mix 2.0 (GeneCopeia, Rockville, MD) and the PCR primers listed in **Table 3**; reactions were carried out using a CFX96 Real Time System (BioRad, Hercules, CA). Each qPCR reaction (10 µl) contained 1.0 µl of template and 200 nM of forward and reverse primers. Thermocycler parameters were: one cycle at 95°C for 10 minutes; followed by 40 cycles of 95°C for 10 seconds, 60°C for 20 seconds, and 72°C for 15 seconds. In each experiment, specificity was confirmed by melting point analyses from 65 °C to 95 °C at 0.5 °C increments. For each transcript and experimental condition, fold change in mutant versus wild-type samples was calculated using the ΔΔCt method normalized to *cam1* as the internal control (48,49). Calculations used the equation: ΔΔCt = [(Ct gene – Ct ref) in wild-type] - [(Ct gene – Ct ref) in mutant], where gene refers to transcript being measured (*atf1, pcr1, php2, php3, php5, rst2*), and ref is the *cam1* reference transcript. Frequencies reported in the figures are mean ± SEM of values from four independent biological replicates.

**Table 3.**
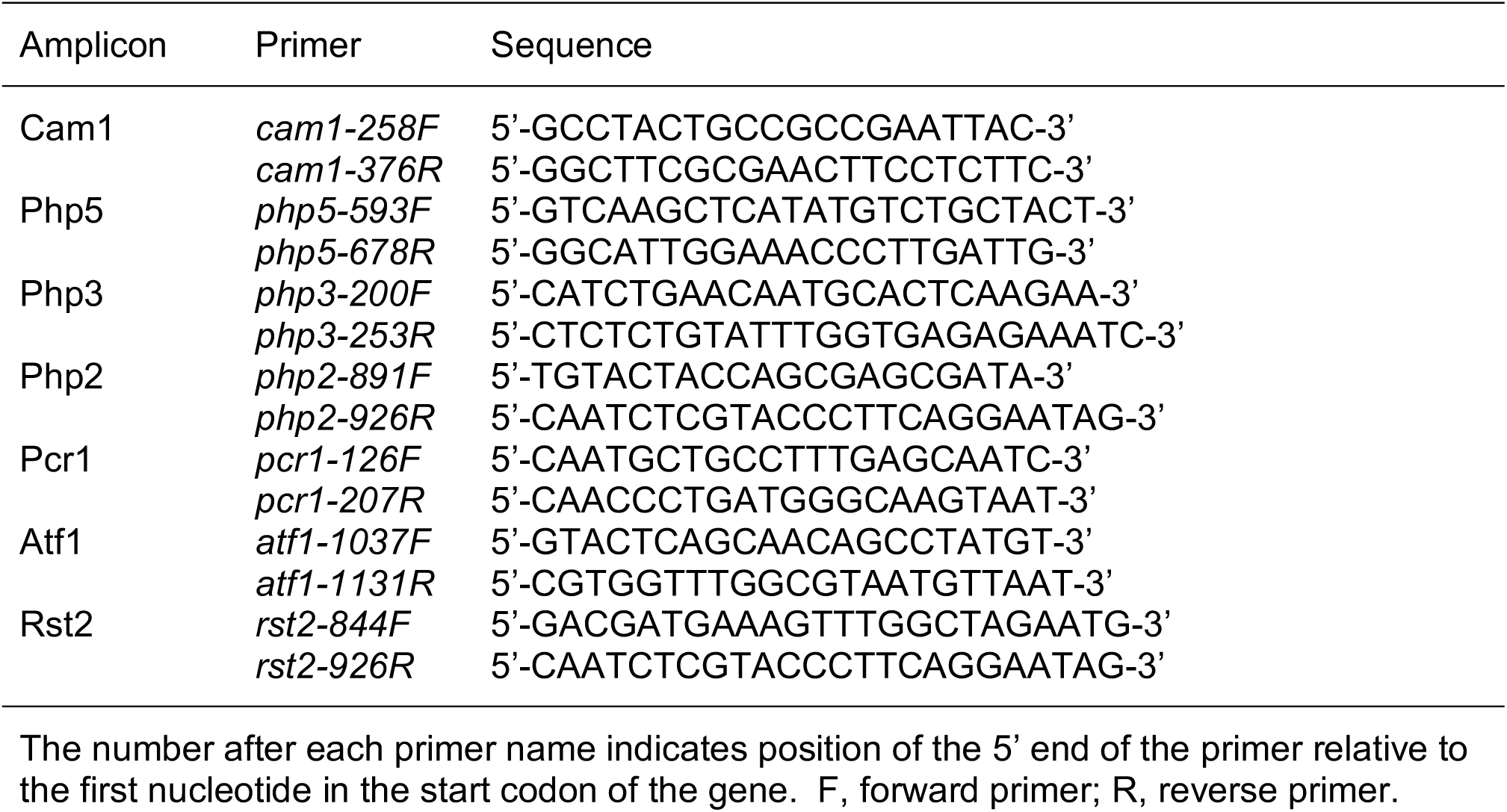
DNA sequences and locations of PCR primers. [upon acceptance for publication, this table will be moved to a supplementary online materials file]

### Replicates and statistics

Each biological replicate included matching controls (e.g., *atf1Δ* versus *atf1*^*+*^), analyzed in parallel under identical conditions, to permit direct, pair-wise comparisons of data between replicates. Data sets were analyzed using paired t-test; differences between means with *p* ≤0.05 were judged to be significant statistically. Linear least squares regression was used to determine the significance of dose-dependent responses.

### Data availability

The authors state that all data necessary for confirming the conclusions presented in the article are represented fully within the article. Yeast strains and other materials generated by the study are available upon request.

## Results

Given that recombination hotspots of fission yeast are activated in *cis* by the binding of transcription factors to their own DNA sites in the genome (10,34,41), we reasoned that cross-talk between transcription factor networks might regulate hotspots indirectly in *trans*. We tested this hypothesis, first, by asking if hotspot-activating transcription factors can regulate DNA sequence-dependent hotspots to which they do not bind. Second, we tested whether the proteins are rate-limiting for promoting recombination at their own DNA binding sites. And third, we tested whether a given hotspot-activating transcription factor can regulate the expression of other hotspot-activating transcription factors. The answer, in each case, is yes.

### Hotspot-activating proteins can regulate hotspot activity of DNA sites to which they do not bind

We compared rates of meiotic recombination in strains bearing either a basal recombination control allele (*M375*) or one of three different hotspot DNA sequence motifs (*M26, CCAAT, Oligo-C*) located near the 5’ end of the *ade6* gene (**Table 1**). Haploid strains with these alleles, which are auxotrophic for adenine, were crossed to another adenine auxotroph that harbors a tester allele (*M210*) near the 3’ end of *ade6* (**Figure 1A**). After mating and meiosis, spores were plated and spore colonies were scored for the frequency of Ade^+^ recombinants. In each case, the hotspot DNA sequence motifs promoted the rate of meiotic recombination substantially (by 900% to 1,800%), relative to that of the basal recombination control (*M375*) (**Figure 1B**).Removing either subunit of the Atf1-Pcr1 heterodimer, which binds to the *M26* DNA site, abolished hotspot activity of *M26* (**Figure 1C**). The rate of recombination fell to that of the basal recombination control allele in wild-type cells. Similarly, subunits of the Php2-Php3-Php5 complex were essential for *CCAAT* motif-promoted recombination (**Figure 1D**) and Rst2 was essential for hotspot activity of its DNA binding site, *Oligo-C* (**Figure 1E**). The results and conclusions are consistent with, and provide independent confirmation of, those reported previously (10,34,40,41): Multiple different *cis*-acting regulatory modules (transcription factors and their DNA binding sites) each position the activity of the basal recombination machinery at hotspots.

**Figure 1.**
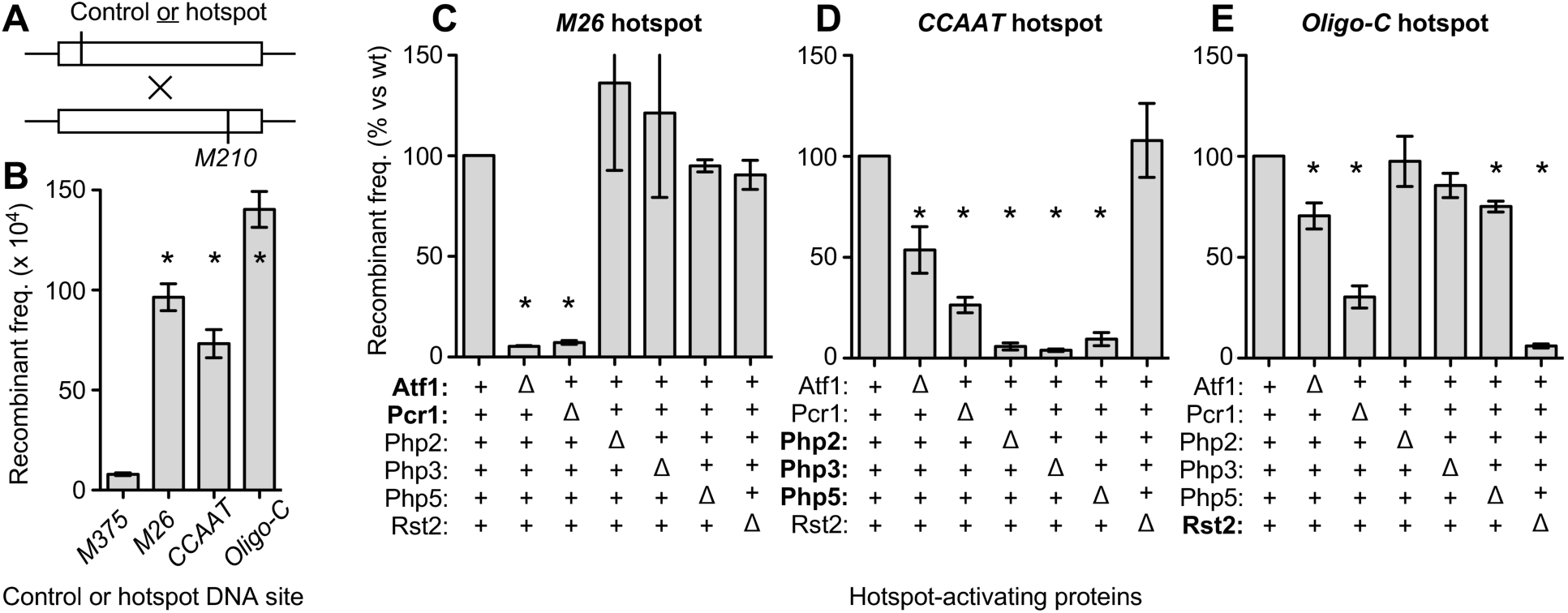
Hotspot-activating proteins can regulate recombination at their own DNA binding sites and at DNA sequence-dependent hotspots to which they do not bind. (**A**) Diagram of genetic crosses showing relative positions of *ade6* alleles (see **Table 1** for sequences of alleles). (**B**) Rates of recombination from crosses bearing hotspot-activating DNA sequence motifs (*M26, CCAAT, Oligo-C*) and a basal recombination control (*M375*) which lacks those DNA sites. (**C-E**) Effects of binding proteins upon DNA sequence-dependent hotspot activity; proteins that bind directly to each hotspot DNA site are highlighted (*bold*). Recombinant frequencies are expressed as percent in null mutant versus wild-type and the results are grouped by class of hotspot: (**C**) *M26* DNA site; (**D**) *CCAAT* DNA site; (**E**) *Oligo-C* DNA site. Data are mean from three or more independent biological replicates; differences with *p* ≤ 0.05 (*) are indicated for hotspot versus control (panel B) and mutant versus wild-type (panels C-E).

When measuring the impact of protein deletions on hotspot activation, we included all possible pair-wise combinations of hotspot-activating proteins and DNA sequence motifs. The removal of the Php2, Php3 or Php5 proteins (which bind to and promote recombination at the *CCAAT* motif) had no significant impact on rates of recombination at the *M26* hotspot (**Figure 1C**). Similarly, the removal of Php2 or Php3 had no significant impact on rates of recombination at the *Oligo-C* hotspot (**Figure 1E**). Likewise, the removal of the Rst2 protein (which binds to and promotes recombination at the *Oligo-C* motif) had no significant impact on the rates of recombination at the *M26* hotspot (**Figure 1C**) or at the *CCAAT* hotspot (**Figure 1D**). These findings suggest that these two DNA sequence-specific, protein-DNA complexes functions with high specificity to promote recombination.

The binding of Atf1-Pcr1 heterodimer to *M26* DNA sites directly activates this class of hotspots (5,34,35). Each subunit of this basic, leucine-zipper (bZIP) heterodimer binds one half-site of the *M26* DNA site (50) and each subunit is essential for hotspot activity (e.g., **Figure 1C**). Strikingly, the removal of Pcr1 significantly reduced (by 74% and 70%, respectively) rates of recombination at the *CCAAT* and *Oligo-C* hotspots (**Figure 1D-1E**). Removing Atf1 also reduced recombination (by 46% and 30%, respectively) for *CCAAT* and *Oligo-C* (**Figure 1D-1E**). The differential effects of Atf1 and Pcr1 on activation of the *CCAAT* and *Oligo-C* hotspots are notable, given that the Atf1-Pcr1 heterodimer regulates *M26*-class hotspots, but these differential effects are readily explained by discovery of underlying molecular mechanisms (described subsequently). [These mechanisms involve the fact that Atf1 and Pcr1 are each multifunctional (e.g., they regulate both recombination and transcription), and they form both homodimers and heterodimers (34,47,50,51)]. Similarly, the removal of Php5 led to a significant, albeit modest reduction in recombination at the *Oligo-C* hotspot (**Figure 1E**). We conclude that hotspot-activating transcription factors can control the activation of heterologous DNA sequence-dependent hotspots to which they do not bind. The most parsimonious model is that transcription factor “A” might regulate the expression level of transcription factor “B”, thereby affecting recombination at “B” DNA sites. This model predicts that DNA sequence motif-dependent hotspots should be sensitive to the abundance of their binding/activating proteins, which we tested as follows.

### Hotspot-activating proteins are rate-limiting for promoting recombination at their own DNA binding sites

To test for dose-dependent responses, we compared rates of recombination for each hotspot DNA sequence motif (*M26, CCAAT, Oligo-C*) in meioses that were homozygous wild-type, heterozygous wild-type/null mutant, and homozygous null mutant for their respective binding proteins. Essentially identical results were observed for each of the six different hotspot-activating proteins (Atf1, Pcr1, Php2, Php3, Php5, Rst2). In every case, the rate of recombination in the heterozygotes was intermediate between that of homozygous wild-type (full hotspot activity) and homozygous null mutant (no hotspot activity) (**Figure 2**). Linear regression analysis of the entire data set revealed a robust, positive correlation between dose and recombination rate (R^2^ = 0.89, *p* < 0.0001). We confirmed this dose-dependent response using a second, independent experimental approach that attenuates the expression of genes without changing their copy numbers (described in a subsequent section). The dose-dependent responses, observed using two different approaches, support a very important conclusion: each hotspot-activating protein is rate-limiting for promoting recombination at its DNA binding site. Consequently, any factor (cellular or environmental) that affects the abundance or functionality of a particular, DNA sequence-specific, hotspot-activating protein will therefore affect rates of recombination at the corresponding class of sequence-dependent hotspots in the genome.

**Figure 2.**
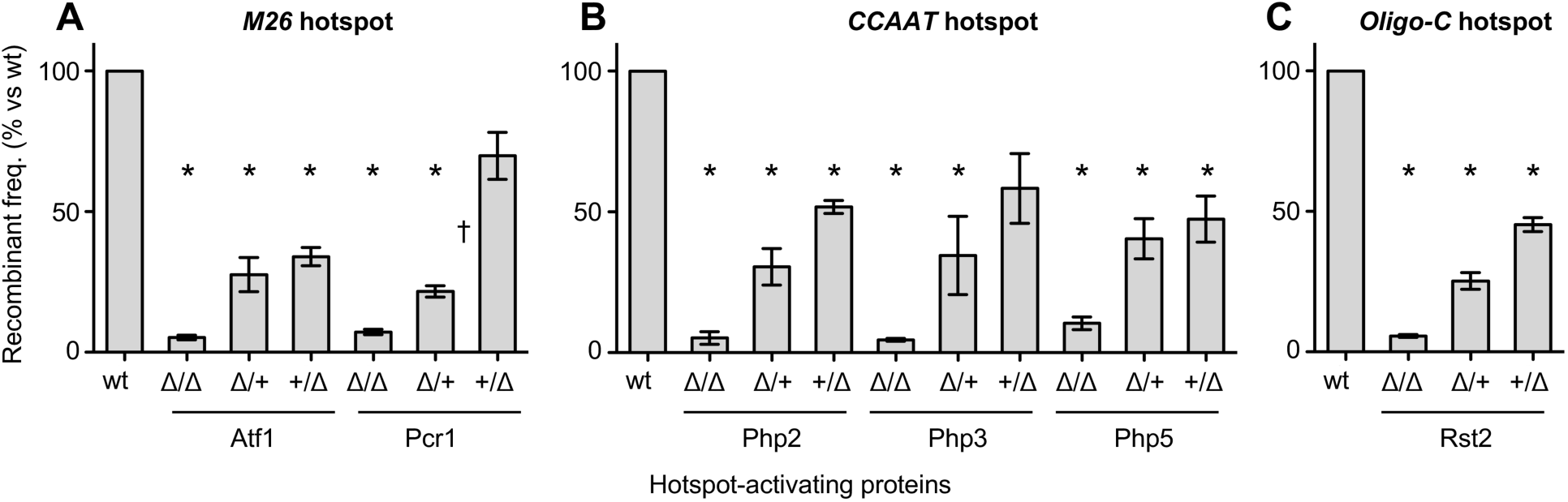
Hotspot activating proteins are rate-limiting and can establish a poised epigenetic state. Recombinant frequencies were determined for crosses that were homozygous wild-type, heterozygous wild-type/null mutant, and homozygous null mutant for hotspot-activating proteins. The heterozygous crosses were in two configurations. In the first configuration (Δ/+), the parent with the hotspot allele was null mutant for the binding protein (e.g., genotype *ade6-M26 atf1Δ*) and the parent with the test-cross allele expressed the binding protein (e.g., *ade6-M210 atf1*^*+*^). In the second configuration (+/Δ), the parent with the hotspot allele was wild-type for the binding protein (e.g., *ade6-M26 atf1*^*+*^) and the parent with the test-cross allele was null mutant for the binding protein (e.g., *ade6-M210 atf1Δ*). Recombinant frequencies are plotted as percent relative to homozygous wild-type and are grouped by class of hotspot: (**A**) *M26* DNA site; (**B**) *CCAAT* DNA site; (**C**) *Oligo-C* DNA site. Data are mean from three or more independent biological replicates; differences with *p* ≤ 0.05 are indicated for mutant versus wild-type (*) and for between heterozygous pairs (†).

### Parental imprinting affects hotspot activation

Analyses of DNA binding in vitro and in vivo (47,51-55), as well as chromatin mapping experiments (29,46,47,51), revealed that the hotspot binding/activating proteins occupy their own DNA binding sites before meiosis. Moreover, the *cis*-acting regulatory modules (sequence-specific protein-DNA complexes) each promote recombination via the modification of chromatin structure before and during meiosis (30). For these reasons, we conducted our analyses of heterozygous wild-type/null mutant meiosis in two reciprocal configurations.

In the first configuration, one haploid parental strain harbored the protein deletion and the hotspot allele (e.g., genotype *atf1Δ ade6-M26*) and the other strain harbored wild-type protein and the tester allele (e.g., *atf1*^*+*^ *ade6-M210*). In the reciprocal configuration, one haploid parental strain harbored wild-type protein and the hotspot allele (e.g., *atf1*^*+*^ *ade6-M26*) and the other strain harbored the protein deletion and the tester allele (e.g., *atf1Δ ade6-M210*). Following mating and during meiosis, each of the two configurations would be heterozygous for the protein (e.g., *atf1*^*+*^*/atf1Δ*) and for the alleles used to measure recombination (e.g., *ade6-M26/ade6-M210*). In other words, in meiosis the genotypes were identical for each paired, reciprocal configuration of protein and DNA site. Therefore, if hotspot activation were simply a function of protein dosage during meiosis, one would expect to observe similar recombinant frequencies from the two reciprocal configurations of each cross. However, the results differed from this expectation.

For each of the six different hotspot binding/activating proteins (Atf1, Pcr1, Php2, Php3, Php5, Rst2), the rate of recombination was higher when the protein was expressed in the haploid parent with the hotspot DNA sequence motif (e.g., *M26*) than when the protein was expressed in the parent with the tester allele (*M210*) (**Figure 2**). This type of parental imprinting suggests that hotspot-regulating protein-DNA complexes can establish a recombinationally poised epigenetic state by the time of karyogamy (i.e., before meiosis). Such poised epigenetic states have important implications for the positioning of recombination throughout the genome and for changes in the distribution of recombination in response to intracellular and extracellular cues (see **Discussion**).

### Hotspot-activating proteins can regulate the expression of other hotspot-activating proteins

Because Atf1 and Pcr1 each controlled the activity of DNA sequence-dependent hotspots to which they do not bind (*CCAAT* and *Oligo-C*) (**Figure 1D-1E**), we tested whether Atf1 and Pcr1 regulate the expression of the proteins that bind to and activate the *CCAAT* box hotspot (Php2-Php3-Php5 complex) and the *Oligo-C* hotspot (Rst2). In fission yeast the depletion of nitrogen source triggers sexual differentiation and the expression of key meiotic drivers (e.g., *ste11*) (51), so we analyzed transcript levels in cells that had been cultured in nitrogen-free media for 60 minutes. Quantitative, real-time, reverse-transcription PCR (qRT-PCR) was used to compare the relative abundance of each mRNA in null mutant cells versus wild-type (**Figure 3**). As expected, no *atf1* transcript was detected in *atf1Δ* mutants and no *pcr1* transcript was detected in *pcr1Δ* mutants.

**Figure 3.**
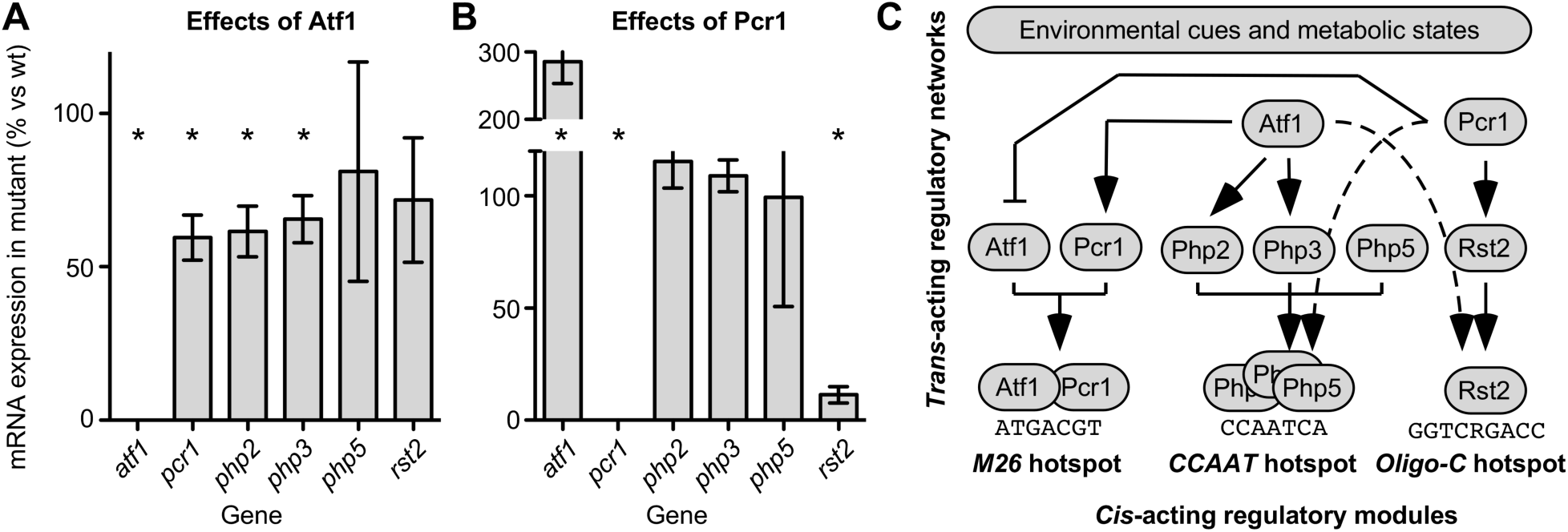
Hotspot-activating proteins can control the expression of proteins that activate other DNA sequence-dependent hotspots. Since Atf1 and Pcr1 regulate hotspots to which they do not bind (**Figure 1**), we tested whether they regulate expression of the other hotspot-activating proteins. Gene expression levels from qRT-PCR in mutants are plotted as percent relative to wild-type for: (**A**) *atf1Δ* mutant; (**B**) *pcr1Δ* mutant. Data are mean from four independent biological replicates; differences with *p* ≤ 0.05 are indicated for mutant versus wild-type (*). (**C**) Conclusions from this and prior figures. Arrows depict statistically significant, linear, dependent pathway relationships elucidated in this study (*solid lines*), as well as connections where significant hotspot control in *trans* was observed, but whose mechanisms do not involve significant changes in expression of the hotspot-activating proteins (*dotted lines*).

It was reported previously, based on quantitative Northern blot analyses of biological samples like those used in this study, that Atf1 promotes the expression of *pcr1* and that Pcr1 represses the expression of *atf1* (47). We obtained the same result using qRT-PCR: the removal of Atf1 led to a reduction in the abundance of the *pcr1* transcript (**Figure 3A**) and the removal of Pcr1 led to an increase in the amount of the *atf1* transcript (**Figure 3B**). The mean relative expression levels that we observed in the mutants versus wild-type (0.6 and 2.9, respectively) were similar to those observed previously (0.5 and 3.4, respectively), which provides reciprocal confirmation of findings in the two different studies and further validates the qRT-PCR approach used in this study.

Overall, considering all six genes analyzed, the removal of Atf1 reduced significantly the expression of *pcr1, php2* and *php3* (**Figure 3A**). The removal of Pcr1 significantly increased the expression of *atf1* and reduced that of *rst2* (**Figure 3B**). We conclude that some hotspot-activating proteins can regulate the expression of other hotspot-activating proteins. We also found that Atf1 had no significant impact on the expression of *php5* and *rst2* (**Figure 3A**) and that Pcr1 did not affect significantly the expression of *php2, php3* and *php5* (**Figure 3B**). Our finding that deletions of *atf1* and *pcr1* affected differentially the expression of the other genes was not surprising, given that Atf1 and Pcr1 (like bZIP proteins in general) form combinatorial homodimers and heterodimers, each of which regulates the expression of a different set of genes (50,53). Nevertheless, the results were highly informative. Collectively these findings— both the positive and negative results for expression-mediated, inter-pathway connections— support important conclusions as to molecular mechanisms.

The finding that Atf1 controls the expression of proteins that bind to and activate the *CCAAT* hotspot (Php2 and Php3; **Figure 3A**), coupled with the fact that those proteins are each rate-limiting for promoting recombination at their own DNA binding site (**Figure 2B**), reveals a specific mechanism by which Atf1 controls in *trans* the activation of the *CCAAT* hotspot (**Figure 1D**). The same conclusion applies for control of the *Oligo-C* hotspot in *trans* by Pcr1 (**Figure 1E**). Pcr1 regulates the expression of Rst2 (**Figure 3B**), which binds to and is rate-limiting for promoting recombination at the *Oligo-C* DNA site (**Figure 2C**). In other words, we documented the same type of fundamental, sequential, transcription-dependent pathway mechanism for the control of two different hotspots (*atf1* → *php2, php3* → recombination at *CCAAT* hotspot; *pcr1* → *rst2* → recombination at *Oligo-C* hotspot; see pathway diagram in **Figure 3C**). This type of signal transduction mechanism, exerted through modulating the expression of rate-limiting activator proteins, has broad implications for how diverse signaling networks can remodel the frequency distribution of recombination across the genome (see **Discussion**).

Interestingly, our qRT-PCR data also indicated that this cannot be the sole mechanism for regulating hotspots in *trans*. Pcr1 was strongly required for the activation of the *CCAAT* box hotspot: rates of recombination in the *pcr1Δ* mutant were only 26% of those in wild-type (**Figure 1D**). However, the removal of Pcr1 had no significant impact on the expression of *php2, php3* and *php5* (**Figure 3B**). The expression levels of these three genes in the *pcr1Δ* mutant were 115%, 109% and 99% of those in wild-type cells. These findings are inconsistent with a model in which the Pcr1-dependent control of this hotspot is solely exerted via altering the expression of its binding/activating proteins, Php2, Php3 and Php5. We infer that there must be at least one additional mechanism by which hotspot-activating transcription factors can control in *trans* other classes of DNA sequence-dependent hotspots (see **Discussion**).

The modulation of expression levels by mutating other factors was also informative for a third reason. In each case where expression of the hotspot-binding/activating protein was reduced (**Figure 3**), there was a corresponding reduction in rates of recombination at its DNA binding site (**Figure 1**). This provides independent support for the conclusion, reported above using a different experimental approach (**Figure 2**), that the hotspot-binding/activating proteins are rate-limiting for promoting recombination at their own DNA binding sites.

## Discussion

In diverse taxa, environmental cues and metabolic states can reshape the frequency distribution of meiotic recombination across the genome (see **Introduction**). We reasoned that previously unknown mechanisms for this plasticity must be related to the way that cells position recombination at hotspots. Therefore, this study analyzed all six of the fission yeast proteins known to occupy, and promote meiotic recombination directly at, their own DNA binding sites (34,41,42). Nearly 200 additional, distinct DNA sequence elements also activate recombination hotspots in fission yeast, but their binding proteins are unknown (40). The mechanisms for modulating the activities of hotspots uncovered in this study likely apply to many, if not most of those other DNA sequence-dependent hotspots. Moreover, because the regulation of hotspots by specific DNA sites and their binding proteins is conserved between species (10,56,57) whose latest common ancestor occurred about 400 million years ago (58), the mechanisms revealed by this study likely apply broadly across taxa. Additional evidence for conservation of mechanisms is described below.

Our main, overall finding—that the activity of each class of DNA sequence-dependent hotspot can be modulated independently in *cis* and in *trans*—is sufficient to support a model for plasticity in the frequency distribution of meiotic recombination (**Figure 4**). Additional evidence that supports this model, its implications, and caveats are described in detail below.

**Figure 4.**
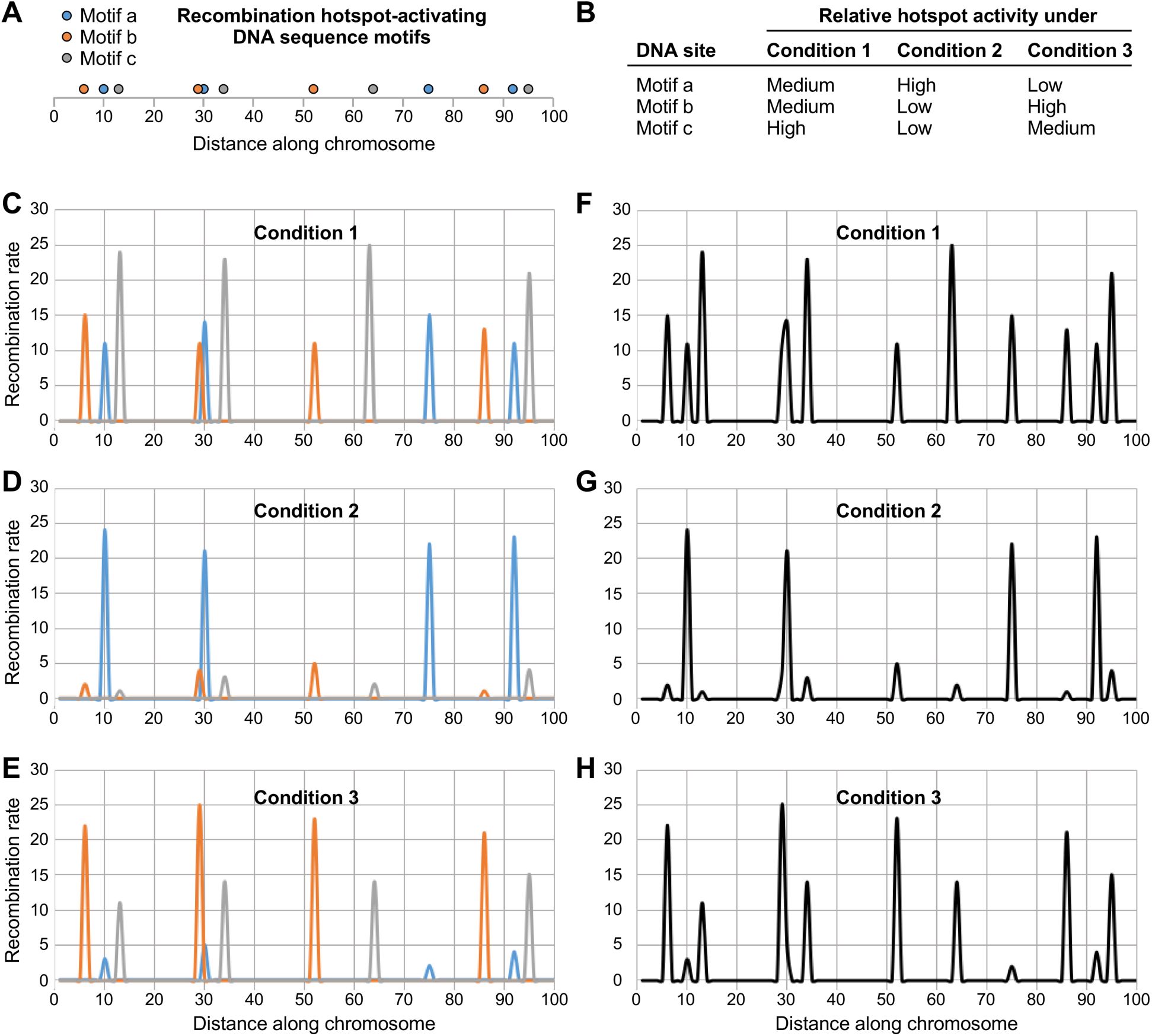
Model for plasticity in the positioning of meiotic recombination. (**A**) Conceptual distribution of three different types of hotspot-activating DNA sequence motifs (*a, blue; b, orange; c, grey*) along the chromosome. (**B**) Each class of DNA sequence-dependent hotspot can be regulated independently, through its own binding/activating proteins, in response to changes in environmental and metabolic conditions. The indicated activities (*low, medium, high*) under each condition were assigned arbitrarily for the sake of illustration. (**C-E**) Contribution of each DNA site to the rate of recombination in its vicinity (*blue, orange, grey lines*) under the experimental conditions listed in panel B. Intensity values of each peak were assigned randomly for low (*1-5*), medium (*11-15*) and high (*21-25*) hotspot activity. (**F-H**) Net impact of the three classes of regulatory DNA sites on the overall distribution of recombination (*black lines*) under the three different conditions. Note differences in the frequency distribution of recombination events. Also note that if one applies a cutoff value for what is a hotspot (e.g., local recombination rate ≤ 10), as is often done in the literature, the number of annotated hotspots and their positions change substantially from condition to condition.

### A model for environmentally and metabolically induced plasticity in the distribution of meiotic recombination

We found that all six of the proteins known to directly regulate meiotic recombination hotspots in fission yeast are rate-limiting for promoting recombination at their own DNA binding sites (**Figure 2**). This property has also been reported for DNA sequence-specific, hotspot activating proteins Rap1 of budding yeast (59) and Prdm9 of mammals (at least in some genetic contexts) (9). Consequently, any signals that affect the functions of a given hotspot binding/activating protein will affect the rate at which it promotes recombination at its own DNA binding sites (conferring none, intermediate or high levels of hotspot activity). Each class of hotspots could be controlled independently, both positively or negatively, by a large network of signals.

The activation of *M26*-class hotspots by the binding of Atf1-Pcr1 heterodimer to *M26* DNA sites (5,34) provides a good example of such multifactorial regulation. Atf1 and Pcr1 are bZIP proteins that can function as homodimers or heterodimers that bind to *M26* or *M26*-like DNA sites (50,51). As transcription factors, these proteins are key responders/effectors of multiple, core environmental stress responses, including those elicited by high temperature (53,60-63). Given that signals of elevated temperature are transduced via promoter-bound Atf1 and Pcr1 to induce the transcription of target genes, and that Atf1-Pcr1-*M26* protein-DNA complexes also promote locally meiotic recombination, one might expect an increase in temperature to enhance rates of recombination at this class of hotspots. Evidence for this can be found in the data of two, prior, independent reports. First, an increase in temperature (from 25 °C to 34 °C) increases the abundance of recombination-initiating DSBs at Atf1-Pcr1-*M26* protein-DNA complex-dependent hotspots (23). Second, for this class of hotspots, there is a strong positive correlation between temperature (spanning six different temperatures from 11 °C to 33 °C) and rates of recombination, as measured in genetic crosses (33).

Atf1 and/or Pcr1 are key responders/effectors of many other signals, including those induced by osmolarity, oxidative state, heavy metals, abundance of nutrients, carbon source, DNA damage and chronological aging (42,47,51,54,60,64-80). These multiple, diverse signals—like those generated by differences in temperature—probably impinge upon Atf1 and Pcr1 to control rates of recombination at *M26*-class hotspots in the genome. Indeed, chromosome-bound Atf1-Pcr1 heterodimer promotes locally both transcription and meiotic recombination via the same general mechanism (chromatin remodeling) (30,51,81,82). This protein-DNA complex can help to seed both euchromatin and heterochromatin (51,83,84). In addition, complex-dependent hotspot activation is regulated both agonistically and antagonistically by multiple different signaling pathways, including those that respond to environmental and metabolic cues (29,46,47,81,85-89). Furthermore, distinct domains of chromosome-bound Atf1-Pcr1 heterodimer promote and repress recombination (42,46), demonstrating that both positive and negative regulation can be exerted directly through the same protein-DNA complex. It thus seems clear that the Atf1-Pcr1 heterodimer directly modulates rates of recombination at *M26*-class hotspots in response to a wide array of different extracellular and intracellular cues.

The same principles for the integration of multiple signals also apply for the *CCAAT* and *Oligo-C* classes of hotspots, which are activated in *cis* by their own transcription factors (Php2-Php3-Php5 complex and Rst2, respectively) (41). Presumably, similar mechanisms extend to at least a subset of the nearly 200 other, short, distinct DNA sequences that activate meiotic recombination hotspots in fission yeast, but whose binding proteins are not yet known (40). The constellation of agonistic and antagonistic signals that affect each of the rate-limiting, hotspot-activating proteins will modulate rates of recombination at their own class of DNA sequence-dependent hotspots. We posit that the independent modulation of recombination rates by each class of DNA sequence-dependent hotspot provides a molecular basis for the previously observed, dynamic changes in the frequency distributions of meiotic recombination that are elicited by intracellular and extracellular cues (e.g., sex or mating type, auxotrophies, nutritional states, temperature; see **Introduction** and references therein). Key features of this new model are depicted in **Figure 4**. We emphasize that the model is a working hypothesis that best explains our new data and many published findings; we do not mean to imply that this model applies universally to all hotspots in all systems.

### Multiple mechanisms for modulating recombination rates via hotspot-activating proteins

One prediction of our model is that hotspot-activating proteins (which are key responders/effectors of environmental and metabolic cues) can regulate the activity of heterologous, DNA sequence-dependent hotspots to which they do not bind. This prediction was confirmed in our study. Atf1 and Pcr1 each controlled rates of recombination at the *CCAAT* and *Oligo-C* hotspots (**Figure 1D-1E**), which are, respectively, bound and activated by the Php2-Php3-Php5 complex and Rst2 (41). Such regulation in *trans* of DNA sequence-dependent hotspots likely applies to other species. For example, in mice the sequence-specific, hotspot-activating protein Prdm9 represses the activity of hotspots to which it does not bind (36,90), although it is not yet known whether those other hotspots are controlled in *cis* by their own DNA sites. Similarly, transcription factors Bas1 and Ino4 of budding yeast affect the frequency distribution of DSBs for hotspots both at their DNA binding sites and elsewhere (37,38), although those studies did not test for DNA sequence dependence of the presumptively direct or indirect effects. In contrast, this study revealed dependent pathway relationships between multiple different proteins and DNA sites (**Figure 1**). Moreover, we identified underlying molecular mechanisms for inter-pathway connections (see pathway diagram in **Figure 3C**). These and additional mechanisms for regulating hotspots are relevant to how hotspots could be controlled by multiple signal transduction networks, not just the discrete inter-pathway examples documented in this study.

The first type of mechanism for controlling hotspot activity is exerted through controlling the abundance of the activator proteins. The exemplar here is that transcription factor “A” can regulate the expression of transcription factor “B” (**Figure 3**), which is rate-limiting for hotspot activity at “B” DNA sites (**Figure 2**). We documented the use of this same type of fundamental, sequential, transcription-dependent pathway mechanism for two different classes of hotspots (*atf1* → *php2, php3* → recombination at *CCAAT* hotspot; *pcr1* → *rst2* → recombination at *Oligo-C* hotspot; see pathway diagram in **Figure 3C**). We note that any other factors (cellular or environmental) whose signals modulate the expression of the rate-limiting, hotspot-binding/activating proteins would also modulate rates of recombination at their DNA sites of action. We do not mean to imply that the regulation of hotspots is exclusively mediated by altering the expression of hotspot-activating proteins. The fact that the activator proteins are rate-limiting supports additional modes of control.

A second type of mechanism for controlling hotspot activity can be exerted via changing the functionality of hotspot-binding/activating proteins without necessarily changing their abundance. Our finding that Pcr1 strongly controls activation of the *CCAAT* box hotspot (**Figure 1D**) without affecting significantly the expression of *php2, php3* or *php5* (**Figure 3B**) supports this idea. There are several ways to achieve robust, transcription-independent control of hotspots in *trans*. For example, “A” could regulate the subcellular localization, DNA binding affinity, or specific activity of “B”, thereby controlling “B”-type hotspots. An excellent case in point is the hotspot-activating Atf1-Pcr1 heterodimer, whose functions are controlled by each of these three different post-translational processes (42,46,47,69,91).

A third, currently hypothetical type of mechanism for controlling hotspot activity lies in the fact that each of the *cis*-acting regulatory modules analyzed in this study promotes recombination through downstream chromatin remodeling pathways (30). If factor “A” controls chromatin remodelers that help to activate “B”-type hotspots, this would contribute to the regulation in *trans*. A recent, proteomics-based approach revealed that many different chromatin remodeling enzymes and histone PTMs function in concert to activate individual hotspots (29), so there are many potential targets for the modulation of hotspot activity at this level.

In summary, there are many classes of DNA sequence-dependent hotspots and multiple different ways to modulate rates of recombination at hotspots via their rate-limiting, hotspot-binding/activating proteins. Each of these known and hypothetical mechanisms provides a nexus for the control of hotspot activity by other factors and signaling networks. As in the Butterfly Effect, minor perturbations to the biological system can, through the diversity of hotspots and regulatory networks, have large impacts on the overall distribution of recombination. Notably, the redistribution of recombination by such mechanisms does not require any changes in the overall distribution of the regulatory DNA sites themselves (e.g., **Figure 4**). Thus, the underlying mechanisms have implications for differences in hotspot usage among taxa and even isolated populations of the same species [92], as well as in response to changing environmental conditions.

### Poised epigenetic states designate the positions of recombination hotspots

As a general principle from species tested, hotspot activation involves the modification of histones and the displacement of nucleosomes, which help localize activity of the basal recombination machinery to the hotspot (1-3). The parental imprinting that we discovered using reciprocally configured, but genetically identical, heterozygous meioses (**Figure 2**) revealed that sequence-specific, hotspot-activating protein-DNA complexes can establish a recombinationally poised epigenetic state prior to meiosis. Recent, temporal analyses of hotspot-recruited chromatin remodeling enzymes and histone PTMs (29), and of chromatin structure at single-nucleosome resolution (30), each support independently this conclusion for the hotspots examined in this study. Some hotspot-specific chromatin remodeling occurs before premeiotic S-phase, further remodeling occurs later in meiosis, leading subsequently to the induction of DSBs. Strikingly, the parental imprinting documented in this study can only be explained by the establishment of hotspot-specific, recombinationally poised epigenetic states before the time of karyogamy between the two haploid parents. This makes the poised epigenetic states the earliest-acting “designators” for the positioning of meiotic recombination defined to date in any organism, which has several important implications.

First, the new and recently published findings (above) indicate that discrete, hotspot-activating DNA sites and their binding proteins are the primary designators temporally, as well as spatially (10), of recombination events in meiosis. This makes sense given that each hotspot-activating protein analyzed in this study is expressed and binds to its DNA sites in vegetative cells, as well as in meiosis (47,51-53,55).

Second, the establishment of poised epigenetic states by hotspot-activating protein-DNA complexes can explain why most hotspot locations already have open chromatin (as judged by nuclease sensitivity assays) before meiosis (28). Given that the activator proteins occupy their DNA sites during vegetative growth, one might expect them to be displaced during, and re-bind following, premeiotic S-phase. Indeed, this dynamic unloading and reloading around the time of premeiotic S-phase has been reported for the Atf1-Pcr1 heterodimer (29). Both the stability of these protein-DNA complexes before meiosis and their rapid reassembly after displacement are presumably driven by the exceptionally high affinity (Kd ∼1 nM) of the Atf1-Pcr1 heterodimer for its DNA binding site, *M26* (50). Our genetic data support such a model (i.e., reloading after displacement in premeiotic S-phase) for each of the hotspots analyzed because we still observed some hotspot activity (albeit to lower levels) when different parents contributed the hotspot DNA site and its binding/activating protein(s) (**Figure 2**). Hence, the new and published findings, when considered together, provide key insight into the process. They indicate that the binding of activator proteins to their DNA sites before karyogamy and their re-binding after displacement in premeiotic S-phase each contribute to chromatin remodeling and hotspot activation.

Third, the poised epigenetic states provide a potential mechanism for the observation that prior environmental conditions can affect the frequency distribution of recombination in subsequent meiosis (27), although at present we cannot tell if the genetic imprinting that we observed extends over multiple, mitotic cell divisions.

## Conclusions

The striking plasticity in the frequency distribution of meiotic recombination across genomes (see **Introduction**) is readily explained by the fact that many, distinct DNA sequence motifs each control rates of recombination locally (see model in **Figure 4**). Each of the sequence-specific DNA binding proteins known to activate meiotic recombination hotspots is rate-limiting for promoting recombination at its own DNA sites. Consequently, each protein or protein complex can respond to agonistic and antagonistic signals to modulate rates of recombination at its own DNA sites. Together, the independent modulation of recombination rates by each class of DNA sequence-dependent hotspot can remodel dynamically the distribution of recombination across the genome in response to environmental and metabolic conditions.

## Acknowledgments

We thank Giulia Baldini, Robert Eoff, Galina Glazko, Aaron Storey, and Alan Tackett for discussions.

